# Discovery of Eremiobacterota with *nifH* homologs in tundra soil

**DOI:** 10.1101/2023.06.30.547195

**Authors:** Igor S. Pessi, Tom O. Delmont, Jonathan P. Zehr, Jenni Hultman

## Abstract

We describe the genome of an Eremiobacterota population from tundra soil that contains the minimal set of *nif* genes needed to fix atmospheric N_2_. This putative diazotroph population, which we name *Candidatus* Lamibacter sapmiensis, links for the first time Eremiobacterota and N_2_ fixation. The integrity of the genome and its *nif* genes are well supported by both environmental and taxonomic signals. *Ca*. Lamibacter sapmiensis contains three *nifH* homologs and the complementary set of *nifDKENB* genes that are needed to assemble a functional nitrogenase. The putative diazotrophic role of *Ca*. Lamibacter sapmiensis is supported by the presence of genes that regulate N_2_ fixation and other genes involved in downstream processes such as ammonia assimilation. Similar to other Eremiobacterota, *Ca*. Lamibacter sapmiensis encodes the potential for atmospheric chemosynthesis via CO_2_ fixation coupled with H_2_ and CO oxidation. Interestingly, the presence of a N_2_O reductase indicates that this population could play a role as a N_2_O sink in tundra soils. Due to the lack of activity data, it remains uncertain if *Ca*. Lamibacter sapmiensis is able to assemble a functional nitrogenase and participate in N_2_ fixation. Confirmation of this ability would be a testament to the great metabolic versatility of Eremiobacterota, which appears to underlie their ecological success in cold and oligotrophic environments.

## Main

The conversion of atmospheric N_2_ to ammonia by microbial diazotrophs represents the largest external source of nitrogen in tundra soils [1–3]. Tundra N_2_ fixation rates are generally low but are predicted to increase with the rise in atmospheric temperatures and precipitation levels associated with anthropogenic climate change [2, 4]. Greater inputs from N_2_ fixation coupled with faster rates of organic matter mineralization will increase the pool of bioavailable nitrogen in tundra soils and potentially lead to higher greenhouse gas emissions, particularly N_2_O [4–6]. More knowledge on the microbial drivers of N_2_ fixation is thus essential for a better understanding of the dynamics of greenhouse gas production in tundra soils.

Here, we investigated the diversity of microbial diazotrophs in the tundra by leveraging a catalogue of 796 manually curated metagenome-assembled genomes (MAGs) from soils in Kilpisjärvi, northern Finland [7] (see *Methods* in **Supplementary information**). The Kilpisjärvi MAGs represent populations from different soil ecosystems (from bare soil to water-logged fens) and encompass a high diversity of Archaea and Bacteria [7, 8]. We searched the 796 Kilpisjärvi MAGs using a hidden Markov model of the *nifH* gene, which encodes the dinitrogenase reductase subunit of the nitrogenase enzyme and is the most widely used marker for N_2_ fixation [9–11]. We found *nifH* homologs in 26 MAGs, most of which are affiliated with groups that include known diazotrophs such as Alpha- and Gammaproteobacteria and methanogenic Archaea (see *‘Kilpisjärvi MAGs with nifH homologs’* in **Supplementary information**).

Surprisingly, we identified *nifH* homologs in the MAG KWL-0264 assigned to Eremiobacterota, a phylum for which no diazotrophic activity and/or nitrogenase sequences have been previously reported. The validity of the MAG as a whole and the inclusion of *nifH* homologs are supported by several features including homogeneous GC content, uniform coverage in multiple metagenomes, and clear taxonomical signal for Eremiobacterota across contigs, including those containing the *nifH* homologs (**Fig. 1a**; see also *‘The integrity of the Eremiobacterota MAG KWL-0264’* in **Supplementary information**). Phylogenomic analysis with other publicly available genomes (*n*=302) confirmed the affiliation of KWL-0264 with Eremiobacterota, placing it alongside other MAGs from polar environments in the family Baltobacteraceae (**Fig. 1b**). The phylogenetic placement and low average nucleotide identity (ANI) with other Eremiobacterota genomes indicate that KWL-0264 represents a distinct lineage in this phylum. Given its higher environmental signal in mineratrophic fens compared to upland soils (see *‘Distribution of nifH MAGs’* in **Supplementary information**), we propose the name *Candidatus* Lamibacter sapmiensis (N.L. fem. n. *lama*, a bog or fen; N.L. masc. n. *bacter*, a rod; N.L. masc. n. *Lamibacter*, a rod that grows on bogs or fens; N.L. masc. adj. *sapmiensis*, pertaining to Sápmi, the cultural region inhabited by the Sámi people in Fennoscandia).

**Fig. 1.**
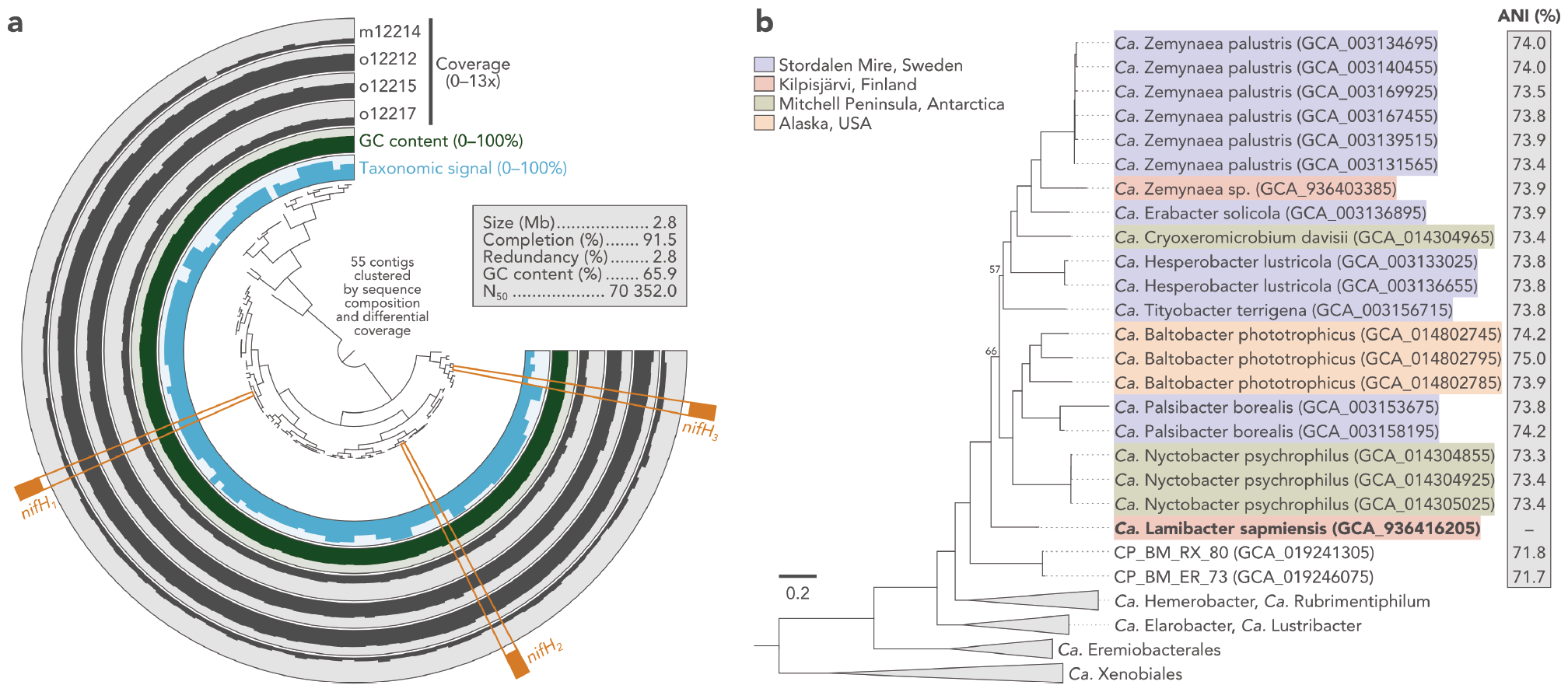
**a)** Representation of the metagenome-assembled genome (MAG) KWL-0264 (*Candidatus* Lamibacter sapmiensis). The four external rings show the mean coverage of each contig across four fen metagenomes from Kilpisjärvi, northern Finland. The innermost blue ring (taxonomic signal) represents the proportion of genes in each contig that had the best match with another Eremiobacterota sequence in the GenBank *nr* database. MAG completion and redundancy was estimated based on the presence of 71 single-copy genes with *anvi’o* v7.1. **b)** Phylogenomic analysis of *Ca*. Lamibacter sapmiensis alongside other publicly available Eremiobacterota genomes (*n*=302). Maximum-likelihood tree (LG+I+G model) based on a concatenated alignment of 71 single-copy genes. Bootstrap support is ≥95% unless shown. The tree is rooted at midpoint. The average nucleotide identity (ANI) between *Ca*. Lamibacter sapmiensis and close neighbours is indicated.

The presence of *nifH* homologs and other related genes suggests that *Ca*. Lamibacter sapmiensis plays a role in N_2_ fixation in tundra soils (**Fig. 2**; see also *‘The nifH homologs of KWL-0264’* in **Supplementary information**). The MAG KWL-0264 contains three *nifH* homologs, which is not uncommon among known diazotrophs [9], and also includes the additional *nifDKENB* genes that are deemed necessary for the assembly of a functional nitrogenase [10, 11]. Importantly, one of the *nifH* homologs (*nifH*_*1*_) encodes a nitrogenase affiliated with Cluster III, which is a diverse group of canonical nitrogenases mostly from anaerobic Bacteria and Archaea with demonstrated diazotrophic activities [9–11]. Further hints towards a role in N_2_ fixation is given by the occurrence of *nifI*_*1*_ and *nifI*_*2*_ downstream of *nifH*_*1*_. These genes are usually found in diazotrophs containing Cluster III nitrogenases and encode a type of GlnB/PII protein that inactivates N_2_ fixation when ammonia is available [9, 12]. It is important to note that the *nifH*_*1*_ homolog is not co-located with *nifDK*, but this is likely due to low genome contiguity rather than gene loss. The other two *nifH* homologs of KWL-0264 (*nifH*_*2*_ and *nifH*_*3*_) encode Cluster IV nitrogenases, which have been largely thought as paralogs that do not participate in N_2_ fixation [9–11]. However, this view has been challenged by the reported diazotrophic growth of *Endomicrobium proavitum* with a Cluster IV nitrogenase [13]. Importantly, the contig with the *nifH*_*2*_ homolog contains the complete set of *nifDKENB* genes, suggesting the capacity to assemble a functional nitrogenase. Further support for a role of *Ca*. Lamibacter sapmiensis in N_2_ fixation is given by the occurrence, near all three *nifH* homologs, of several genes that are involved in nitrogen and amino acid transport and metabolism. These include genes for the GS-GOGAT pathway of ammonia assimilation (*glnA* and *gltD*), ABC-type branched-chain amino acid (*livFGHKM*) and nitrate (*tauABC*) transporters, and amino acid biosynthesis (*dapD, aspB, cysE, argA*, and *metC*).

**Fig. 2.**
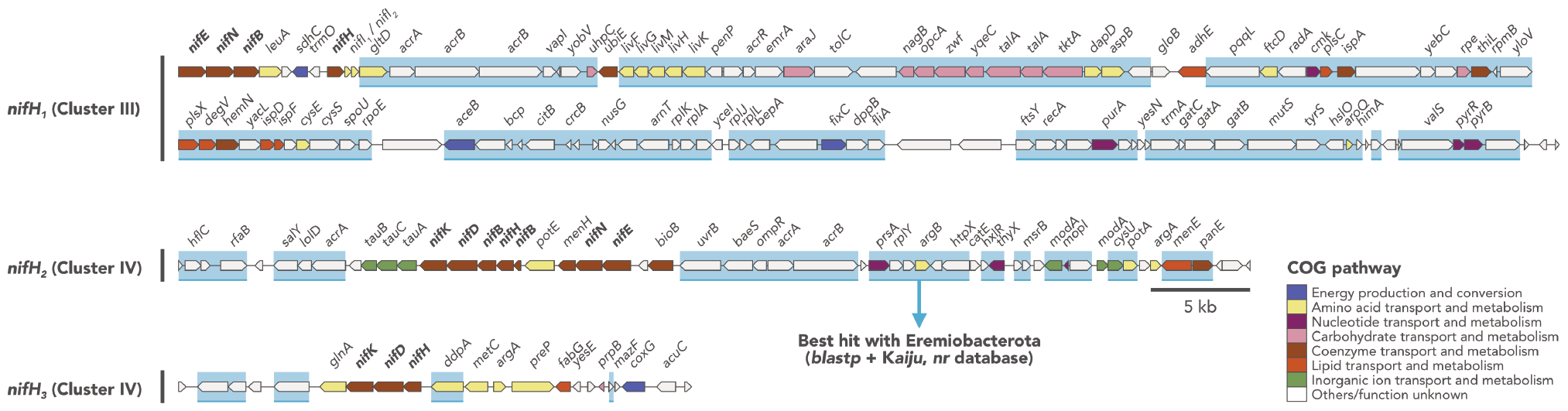
Representation of the three contigs harbouring *nifH* homologs in *Candidatus* Lamibacter sapmiensis KWL-0264. Genes highlighted with a blue rectangle had the best match with another Eremiobacterota sequence in the GenBank *nr* database.

Further gene annotation and metabolic reconstruction revealed many potential traits of *Ca*. Lamibacter sapmiensis that might be beneficial in mineratrophic fens and other oligotrophic environments (**Fig. 3**; see also *‘Reconstruction of the metabolic potential of KWL-0264’* in **Supplementary information**). First, the potential to fix N_2_ represents an ecological advantage in northern fens and peatlands in general, where most of the nitrogen is bound to the organic matter and thus not readily available for growth [3, 5]. Second, KWL-0264 encodes the potential to fix CO_2_ using energy obtained from the oxidation of H_2_ and CO. This metabolic strategy, known as atmospheric chemosynthesis, has been reported for other Eremiobacterota, and appears to underlie their ecological success in cold and oligotrophic ecosystems such as tundra soils, polar deserts, and peatlands [14–17]. Third, the potential to oxidize H_2_ can help alleviate the high energetic cost of N_2_ fixation, as it allows the recovery of part of the spent energy by recycling the H_2_ that is produced during the process [18]. Finally, KWL-0264 encodes a clade II N_2_O reductase and thus the potential for anaerobic respiration via the reduction of N_2_O to N_2_, which is favoured in the typically anoxic environment of water-logged fens [7]. Moreover, it can be speculated that the reduction of N_2_O to N_2_ might also be used as a strategy to fuel N_2_ fixation. The ability to reduce N_2_O indicates a potential role of this Eremiobacterota population in mitigating the emission of this potent greenhouse gas, as has been shown for other non-denitrifying N_2_O reducers [19].

**Fig. 3.**
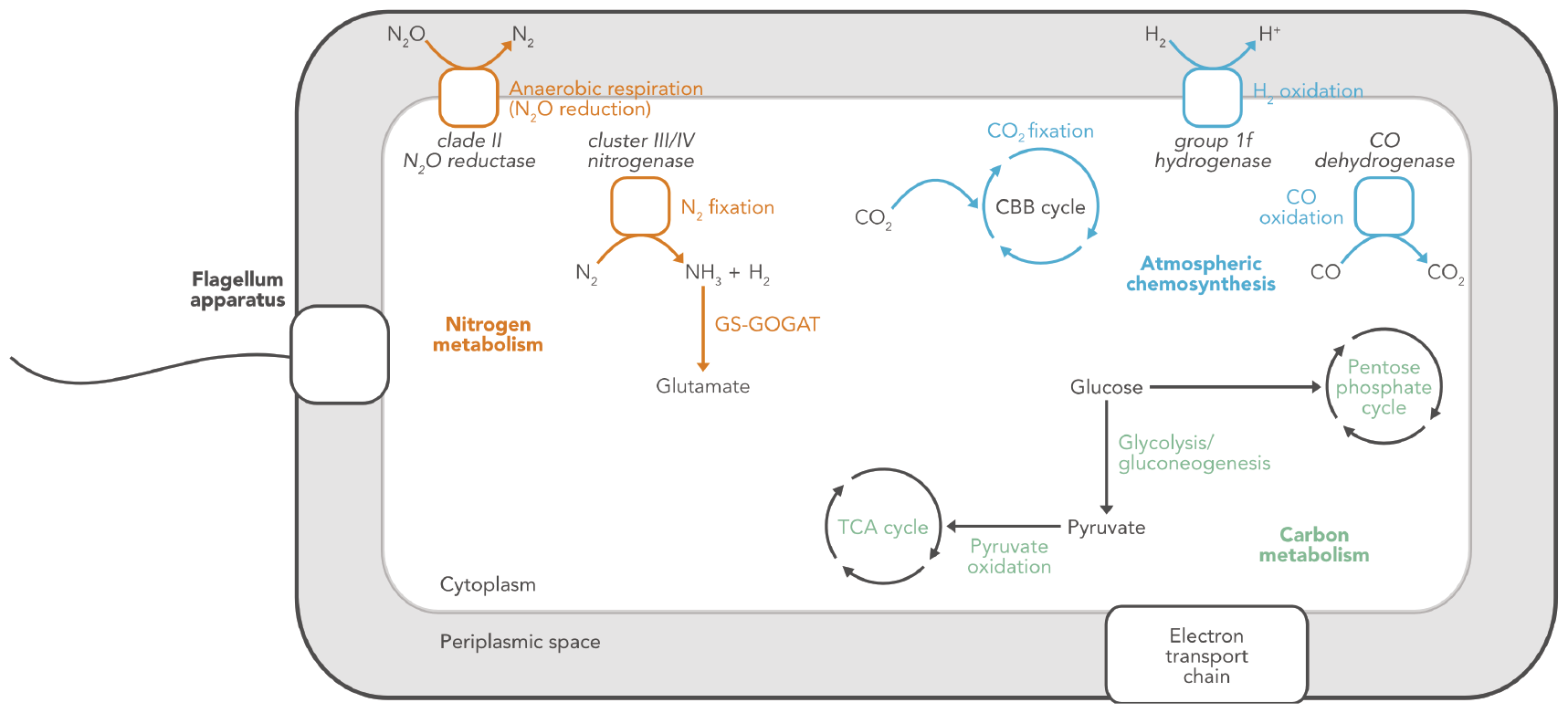
Simplified representation of the metabolic potential of *Ca*. Lamibacter sapmiensis KWL-0264. Only selected pathways discussed in the text are shown. Figure created with [BioRender](https://www.biorender.com/).

After more than 20 years since the first description of Eremiobacterota as “candidate phylum WPS-2” from contaminated soil [20], the recovery of a MAG harbouring *nifH* homologs suggests that N_2_ fixation might also be a part of the metabolic repertoire of this phylum. At present, it remains unclear if the genomic constitution of *Ca*. Lamibacter sapmiensis can lead to a nitrogenase with function in N_2_ fixation. Further studies should focus on isolating this population *in vitro*, recovering other closely related genomes (preferentially using long-read sequencing technologies), and providing evidence of N_2_ fixation activity using, for example, rate measurements coupled with (meta)transcriptomics and/or stable isotope probing.

## Author contributions

**Igor S. Pessi:** conceptualization (supporting), data curation (lead), formal analysis (lead), investigation (lead), methodology (lead), visualization, writing – original draft preparation (lead), writing – review & editing (lead). **Tom Delmont:** methodology (supporting), writing – review & editing (supporting). **Jonathan P. Zehr:** methodology (supporting), writing – review & editing (supporting). **Jenni Hultman:** conceptualization (lead), funding acquisition (lead), methodology (supporting), project administration (lead), writing – review & editing (supporting).

## Acknowledgements

We would like to acknowledge the CSC – IT Centre for Science, Finland for providing the computing resources, and Antti Karkman, Eeva Eronen-Rasimus, and Hermanni Kaartokallio for comments. We also thank the two anonymous reviewers for the critical evaluation of our manuscript. This study was funded by the Academy of Finland (project 335354) and the University of Helsinki.

## Data availability statement

The raw metagenomic data and the 796 Kilpisjärvi MAGs including *Ca*. Lamibacter sapmiensis KWL-0264 can be found in the European Nucleotide Archive (ENA) under the project PRJEB41762. The MAGs are also available from FigShare (doi.org/10.6084/m9.figshare.19722505). The detailed bioinformatics workflow used in this study can be found in github.com/ArcticMicrobialEcology/Candidatus-Lamibacter-sapmiensis.

