## Supplementary information for "Discovery of Eremiobacterota with *nifH* homologs in tundra soil"

### Methods

##### Description of the dataset

The dataset used in this study comprises 796 metagenome-assembled genomes (MAGs) recovered from tundra soils in Kilpisjärvi, northern Finland (Pessi *et al.*, 2022; see also github.com/ArcticMicrobialEcology/Kilpisjarvi-MAGs). In brief, 69 metagenomes were sequenced with Illumina NextSeq/NovaSeq and assembled with *MEGAHIT* v1.1.1.2 (Li *et al.*, 2015). Two additional metagenomes were generated with Nanopore MinION and assembled with *metaFlye* v2.7.1 (Kolmogorov *et al.*, 2020). The assemblies were imported to *anvi'o* v6.2 (Eren *et al.*, 2021) and contigs were manually binned into MAGs based on differential coverage and tetranucleotide frequency using the *anvi-interactive* interface (merenlab.org/2016/02/27/the-anvio-interactive-interface.) Each MAG was manually inspected using the *anvi-refine* interface (merenlab.org/2015/05/11/anvi-refine) to identify and remove contigs with discrepant environmental, compositional, and phylogenetic signals (based on differential coverage, tetranucleotide frequency, and taxonomy of single-copy genes, respectively). The final set of 796 Kilpisjärvi MAGs is available from FigShare (doi.org/10.6084/m9.figshare.19722505) and ENA (ebi.ac.uk/ena/browser/view/PRJEB41762). The latter also includes the raw data for the 69 Illumina and the two Nanopore metagenomes.

##### Analysis of *nifH* homologs

The detailed and reproducible bioinformatics workflow used in this study can be found in github.com/ArcticMicrobialEcology/Candidatus-Lamibacter-sapmiensis. The 796 Kilpisjärvi MAGs were imported to *anvi'o* v7.1 (Eren *et al.*, 2021) and gene calls were identified with *Prodigal* v2.6.3 (Hyatt *et al.*, 2010). *HMMER* v3.3.2 (Eddy, 2011) was then used to search for *nifH* homologs based on the KOfam hidden Markov model (HMM) K02588 applying the pre-computed, domain-specific bit score threshold (Aramaki *et al.*, 2020). Putative *nifH* homologs were further analysed by searching the *nr*, *RefSeq*, and *Swiss-Prot* databases with *blastp* v2.14.0 (Camacho *et al.*, 2009). Phylogenetic analysis of *nifH* homologs was done alongside selected sequences from cultured diazotrophs (obtained from jzehrlab.com/nifh) and the sequences reported by North *et al.* (2020). For this, amino acid sequences were aligned with *MAFFT* v7.520 (Katoh & Standley, 2013), the alignment was trimmed with *trimAl* v1.4 (Capella-Gutiérrez *et al.*, 2009), and a maximum-likelihood tree was computed with *IQ-TREE* v.2.2.2.7 (Nguyen *et al.*, 2015) using the LG+R10 model and 1000 bootstrap replicates according to North *et al.* (2020).

**Analysis of MAGs containing *nifH* homologs**

MAGs with *nifH* homologs were imported to *anvi'o* v7.1 (Eren *et al.*, 2021), in which i) gene calls were identified with *Prodigal* v2.6.3 (Hyatt *et al.*, 2010); ii) *HMMER* v3.3.2 (Eddy, 2011) was used to find a set of 71 bacterial and 76 archaeal single-copy genes (modified from Lee, 2019); iii) *DIAMOND* v2.1.7.161 (Buchfink *et al.*, 2015) was used to assign taxonomy to the single-copy genes according to the Genome Taxonomy Database (GTDB) r95 (Parks *et al.*, 2022); and iv) genome-wide annotation was done against the KOfam database (Aramaki *et al.*, 2020) with *HMMER* v3.3.2 (Eddy, 2011) and the COG database (Galperin *et al.*, 2021) with *DIAMOND* v2.1.7.161 (Buchfink *et al.*, 2015). The MAGs were classified with *GTDB-Tk* v1.5.0 (Chaumeil *et al.*, 2020) and the GTDB r202 (Parks *et al.*, 2022).

**Read recruitment analysis**

Read recruitment was used to estimate the abundance of *nifH* MAGs across the metagenomes from which they originated (Pessi *et al.*, 2022). First, *pyANI* v0.2.12 (Pritchard *et al.*, 2016) was used to compute the pairwise average nucleotide identity (ANI) between the MAGs. This revealed that each MAG represents a unique lineage (maximum pairwise ANI of 84.1%), and thus dereplication was not needed for downstream analyses. The metagenomic reads were filtered and trimmed (minimum Phred score of 20 and minimum length of 50 bp) with *Cutadapt* v1.16 (Martin, 2011) and mapped to the MAGs with *bowtie2* v2.3.5 (Langmead & Salzberg, 2012) and *SAMtools* v1.10 (Li *et al.*, 2009). *CoverM* v0.6.1 (github.com/wwood/CoverM) was then used to summarize the abundance of each MAG across the metagenomes. Alignments with <95% identity and <75% aligned fraction were discarded, and MAG abundances were normalized to reads per kilobase per million reads mapped (RPKM) to account for differences in library and genome size. In *R* v4.2.2 (r-project.org), RPKM values were used to compute the alpha diversity (richness and Shannon index) of each community with the *vegan::diversity()* function (cran.r-project.org/web/ packages/vegan), and differences between the ecosystems were tested using one-way ANOVA with the *stats::lm()* function (stat.ethz.ch/R-manual/R-devel/library/stats/ html/00Index.html). Differences in community composition (beta diversity) between the ecosystems were tested using permutational multivariate ANOVA (PERMANOVA) with the *vegan::adonis2* function (cran.r-project.org/web/packages/vegan) based on Bray-Curtis distances and 999 permutations.

**Analysis of the Eremiobacterota MAG KWL-0264**

Further analyses were carried out for the Eremiobacterota MAG KWL-0264. As another assessment of the integrity of the MAG, a taxonomic label was assigned to each gene call by searching the GenBank *nr* database using *blastp* v2.14.0 (Camacho *et al.*, 2009) and *Kaiju* v1.9.2 (Menzel *et al.*, 2016). A taxonomic signal was then calculated for each contig as the proportion of gene calls that had the best match (highest bitscore) with another Eremiobacterota sequence according to both *blastp* and *Kaiju*. Phylogenomic analysis was done alongside other Eremiobacterota genomes (*n*=302) from GenBank (accessed on 14 November 2023), Ji *et al.* (2021), and Pessi *et al.* (2022). The genomes were retrieved with *ncbi-genome-download* v0.3.3 (github.com/kblin/ncbi-genome-download) and imported to *anvi'o* v7.1 (Eren *et al.*, 2021) as described above for the Kilpisjärvi MAGs. Amino acid sequences for a set of 71 bacterial single-copy genes (modified from Lee, 2019) were then retrieved and aligned with *MAFFT* v7.520 (Katoh & Standley, 2013). The alignments were concatenated and a maximum-likelihood tree was computed with *IQ-TREE* v.2.2.2.7 (Nguyen *et al.*, 2015) using the automated model selection and 1000 bootstrap replicates. The ANI between KWL-0264 and the other Eremiobacterota genomes was estimated with *pyANI* v0.2.12 (Pritchard *et al.*, 2016). Further annotation and metabolic reconstruction of the MAG KWL-0264 was obtained with *DRAM* v1.4.6 (Shaffer *et al.*, 2020) implemented on KBase (Arkin *et al.*, 2018). Key genes involved in atmospheric chemosynthesis (hydrogenase, CO dehydrogenase, and RuBisCO) were submitted to phylogenetic analysis alongside sequences from Søndergaard *et al.* (2016), Cordero *et al.* (2019), and Yabe *et al.* (2022), respectively. The phylogenetic analyses were done as described above for the *nifH* homologs but using the automated model selection. Several attempts were made to improve the contiguity of the MAG KWL-0264, but none resulted in a better (more contiguous) MAG. First, metagenomic assemblies were produced with *metaSpades* v3.15.5 (Nurk *et al.*, 2017) for the three samples where the MAG was detected at highest coverage (samples o12212, o12215, and o12217). Second, the sample o12212 was sequenced with Nanopore MinION and assembled with *metaFlye* v2.9.2 (Kolmogorov *et al.*, 2020), and a hybrid assembly of Illumina and Nanopore data was also produced with *metaSpades* v3.15.5 (Nurk *et al.*, 2017). Finally, *MetaCarvel* v1.1 (Ghurye *et al.*, 2019) and *Binnacle* v1.0 (Muralidharan *et al.*, 2021) were used to scaffold and extend the original contigs of the MAG KWL-0264.

#### Supplementary results

##### Kilpisjärvi MAGs with *nifH* homologs

We identified 29 putative *nifH* homologs distributed across 26 of the 796 Kilpisjärvi MAGs, all of which also contain *nifDK* (**Fig. S1a, Table S1**). Phylogenetic placement based on the GTDB release r202 assigned the 26 MAGs to the phyla Proteobacteria (Alphaproteobacteria, *n*=2; Gammaproteobacteria, *n*=6), Nitrospirota (*n*=5), Methanobacteriota (*n*=4), Halobacteriota (*n*=2), Actinobacteriota (*n*=2), Desulfobacterota (*n*=2), Firmicutes (*n*=1), Myxococcota (*n*=1), and Eremiobacterota (*n*=1). In the NCBI taxonomy (ncbi.nlm.nih.gov/ taxonomy), the phyla Halobacteriota and Methanobacteriota are part of the phylum Euryarchaeota, and the phyla Desulfobacterota and Myxococcota are included in the class Deltaproteobacteria (phylum Proteobacteria). Phylogenetic analysis placed most of the putative *nifH* homologs (*n*=23) alongside canonical nitrogenases from Clusters I, II, and III (**Fig. S1b, Table S1**). The remaining *nifH* homologs (*n*=6) were grouped with Cluster IV nitrogenases. All the 29 *nifH* homologs encode conserved residues that are associated with nitrogenase activity (Zheng *et al.*, 2016; North *et al.*, 2020; Dong *et al.*, 2022): MgATP binding, ATP hydrolysis and [$_4$Fe$_4$S] cluster coordination (Cys$_{97}$ and Cys$_{132}$), and ADP-ribosylation (Arg$_{100}$) (**Fig. S1c**).

One MAG assigned to *Candidatus* Patescibacteria (KWL-0212) was discarded from the dataset because discrepancies in taxonomic and coverage signal across contigs indicate that the MAG is likely a chimeric artifact (**Fig. S2**). Taxonomic assignment of single-copy genes revealed that around half of the KWL-0212 genome (16 contigs, 526 kb) comprises contigs without a taxonomic signal for *Ca.* Patescibacteria, including the contig where the *nifHDK* homologs are located. Furthermore, read recruitment analysis using the metagenomic datasets from which KWL-0212 was assembled (Pessi *et al.*, 2022) revealed that the contigs without a taxonomic signal also display an aberrant coverage profile compared to the contigs assigned to *Ca.* Patescibacteria (four contigs, 666 kb). These discrepancies in taxonomic signal and coverage profile indicate that the MAG KWL-0212 is a chimeric artifact and that the original *Ca.* Patescibacteria population likely does not contain nitrogenase genes. The record of the MAG KWL-0212 in ENA (accession GCA_936414605) was updated to remove the 16 contigs with aberrant taxonomic and coverage signals.

##### Distribution of *nifH* MAGs

Recruitment analysis of metagenomic reads from different tundra soil ecosystems in Kilpisjärvi, northern Finland (Pessi *et al.*, 2022), revealed differences between the *nifH*-containing communities in upland (barren, heathland, and meadow) and fen soils (**Fig. S3**). Alpha diversity estimates were lower in the upland metagenomes compared to the water-logged fens (ANOVA; $R^2 = 0.38–0.80$, $p < 0.001$), and beta diversity analysis showed that community composition also differs significantly (PERMANOVA; $R^2 = 0.30$, $p < 0.0001$). In general, upland metagenomes were dominated by a few MAGs encoding Cluster I nitrogenases (Alphaproteobacteria, Gammaproteobacteria, and Myxococcota), whereas fen communities were more diverse. This mirrors the patterns observed for the whole archaeal and bacterial communities in these sites, in which fens also appeared more diverse than upland soils (Pessi *et al.*, 2022).

#### The integrity of the Eremiobacterota MAG KWL-0264

Several lines of evidence indicate that the Eremiobacterota MAG KWL-0264 is not a binning and/or assembly artifact. First, estimates based on the presence of 71 single-copy core genes indicate that the MAG is 91.5% complete and has a low redundancy level of 2.8% (**Table S1**). Moreover, the MAG has homogeneous GC content and coverage throughout all contigs and across multiple samples (**Fig. S4**). Importantly, most contigs (40 out of 55) have a strong taxonomic signal for Eremiobacterota, including the contigs with the *nifH* homologs. For instance, 99 of the 122 genes in the contig where *nifH₁* is located had their best match in the GenBank *nr* database to a sequence from Eremiobacterota. These included five genes encoding the ribosomal proteins RplA, RplJ, RplK, RplL, and RpmB. Finally, the likelihood of misassembly is low given that the *nif* genes and their flanking genes are bridged by many paired-end reads. Taken together, these observations clearly indicate that the *nifH*-containing MAG KWL-0264 represents a real Eremiobacterota population from tundra soils. We did not find *nifH* homologs in any other publicly available Eremiobacterota genome (*n*=302), which indicates that KWL-0264 is at present the only member of this group that encodes the potential for $N_2$ fixation.

#### The *nifH* homologs of KWL-0264

The MAG KWL-0264 contains three *nifH* homologs (**Fig. S1b**). One of these (*$nifH_1$*) encodes a Cluster III nitrogenase that is related to sequences from Desulfobacterota (e.g. *Desulfovibrio vulgaris* and *Desulfarculus baarsi*) and Verrucomicrobiota (e.g. "Opitutaceae bacterium" and *Coraliomargarita akajimensis*). However, the contig containing the Cluster III *$nifH_1$* homolog lacks *nifDK*, which encode the nitrogenase subunit that contains the active site for $N_2$ fixation and is thus essential for the activity of the enzyme (Zehr *et al.*, 2003; Dos Santos *et al.*, 2012; Koirala & Brözel, 2021). It seems unlikely that a functional Cluster III nitrogenase could be assembled from *$nifH_1$* without *nifDK*. However, given that KWL-0264 does not represent a complete, circular genome, and that the *$nifH_1$* homolog is located towards the start of the contig, it is possible to attribute the lack of adjacent *nifDK* genes to the fragmented nature of the MAG resulting from challenges in metagenomic assembly (Meyer *et al.*, 2022). The other two *nifH* homologs of KWL-0264 (*$nifH_2$* and *$nifH_3$*) are affiliated with Cluster IV nitrogenases and clustered alongside sequences from Alphaproteobacteria (*Rhodopseudomonas*, *Rhodomicrobium*, and *Rhodospirillum*) and Euryarchaeota (*Methanoregula* and *Methanosphaerula*), respectively. The complete set of *nifDKENB* genes is found in the contig containing the *$nifH_2$* homolog, while *$nifH_3$* is co-located with *nifDK* but not *nifENB*. In addition to a potential role in $N_2$ fixation of Cluster IV nitrogenases (Zheng *et al.*, 2016), it is possible that the *nifDK* genes located downstream of *$nifH_2$* and *$nifH_3$* can compensate for the lack of these genes near the Cluster III *$nifH_1$* homolog. However, this would require the co-expression of genes that are potentially located far apart from each other, the likelihood of which is difficult to assess at present without a better spatial resolution of the genomic organisation of KWL-0264.

**Reconstruction of the metabolic potential of KWL-0264**

Gene annotation and metabolic reconstruction revealed that KWL-0264 encodes the potential for atmospheric chemosynthesis, consisting of $CO_2$ fixation via the Calvin-Benson-Bassham (CBB) cycle with a type 1e RuBisCo, $H_2$ oxidation with a high-affinity group 1f Ni-Fe hydrogenase, and CO oxidation with a CO dehydrogenase (**Fig. S5**, **Table S2**). Interestingly, KWL-0264 encodes a clade II $N_2O$ reductase (*nosZ*) and thus the potential for anaerobic respiration via $N_2O$ reduction (**Table S2**). In addition, KWL-0264 encodes genes for the GS-GOGAT pathway of ammonia assimilation (*e.g. glnA*, glutamine synthetase; *gltD*, glutamate synthase), the core module of glycolysis/gluconeogenesis (*e.g. gap*, glyceraldehyde phosphate dehydrogenase; *eno*, enolase), pyruvate oxidation (*e.g. pdh*, pyruvate dehydrogenase), the tricarboxylic acid (TCA) cycle (*e.g. idh*, isocitrate dehydrogenase; *suc*, succinyl-CoA synthetase), the pentose phosphate cycle (*e.g.* G6PD, glucose-6-phosphate dehydrogenase; PGD, 6-phosphogluconate dehydrogenase), the complexes I–V of the electron transport chain (*e.g. nuo*, NADH-quinone oxidoreductase; *sdh*, succinate dehydrogenase; *cox*, cytochrome c oxidase; ATPF, F-type ATPase), and the flagellum apparatus (*e.g. fliF*, flagellar M-ring protein; *fliE*, flagellar hook protein). KWL-0264 does not have any genes involved in the phosphotransferase system (PTS) of carbohydrate uptake and does not encode any carbohydrate-active enzyme (CAZy). Unlike other Eremiobacterota, KWL-0264 does not encode the potential for anoxygenic photosynthesis (*puf* and *bch* genes).

### Supplementary tables and figures

**Table S1.** Information on 26 metagenome-assembled genomes (MAGs) containing *nifH* homologs recovered from tundra soils in Kilpisjärvi, northern Finland.

| MAG | Classification [1] | *nifH* cluster | Size (Mb) | # contigs | GC (%) | Compl. (%) [2] | Redund. (%) [2] |
|---|---|---|---|---|---|---|---|
| KWL-0043 | Halobacteriota; *Methanosarcina* | III | 3.2 | 667 | 37.8 | 59.2 | 5.3 |
| KWL-0479 | Halobacteriota; *Methanoregula* | III, IV | 1.3 | 261 | 49.7 | 50.0 | 2.6 |
| KWL-0006 | Methanobacteriota; *Methanobacterium* | IV | 1.0 | 253 | 34.6 | 63.2 | 5.3 |
| KWL-0003 | Methanobacteriota; *Methanobacterium* | IV | 1.6 | 311 | 35.1 | 68.4 | 5.3 |
| KWL-0017 | Methanobacteriota; *Methanobacterium* | II | 1.2 | 67 | 36.3 | 53.9 | 0.0 |
| KWL-0007 | Methanobacteriota; *Methanobacterium* | II | 1.1 | 97 | 36.1 | 56.6 | 0.0 |
| KWL-0579 | Actinobacteriota; Solirubrobacteraceae | III | 2.7 | 508 | 66.7 | 53.5 | 4.2 |
| KWL-0387 | Actinobacteriota; *Demequina* | III | 3.2 | 694 | 65.8 | 69.0 | 2.8 |
| KWL-0264 | Eremiobacterota; Baltobacteraceae | III, IV | 2.8 | 55 | 65.9 | 91.5 | 2.8 |
| KWL-0436 | Firmicutes; Clostridiaceae | III | 2.4 | 159 | 36.1 | 54.9 | 1.4 |
| KWL-0503 | Desulfobacterota; Smithellaceae | IV | 1.2 | 291 | 44.0 | 71.8 | 5.6 |
| KWL-0475 | Desulfobacterota; Desulfomonilaceae | III | 2.3 | 411 | 46.5 | 73.2 | 4.2 |
| KWL-0282 | Nitrospirota; Thermodesulfovibrionales | I | 3.4 | 119 | 47.6 | 94.4 | 1.4 |
| KWL-0284 | Nitrospirota; Thermodesulfovibrionales | I | 1.8 | 388 | 49.9 | 83.1 | 5.6 |
| KWL-0205 | Nitrospirota | I | 1.8 | 32 | 54.0 | 81.7 | 2.8 |
| KWL-0286 | Nitrospirota | I | 3.3 | 56 | 58.1 | 60.6 | 4.2 |
| KWL-0173 | Nitrospirota | I | 2.1 | 331 | 53.9 | 62.0 | 1.4 |
| KWL-0075 | Myxococcota; Anaeromyxobacteraceae | I | 2.3 | 306 | 71.6 | 50.7 | 1.4 |
| KWL-0169 | Alphaproteobacteria; Xanthobacteraceae | I | 2.3 | 220 | 64.8 | 62.0 | 0.0 |
| KWL-0235 | Alphaproteobacteria; *Methylocella* | I | 1.8 | 308 | 58.9 | 71.8 | 4.2 |
| KWL-0288 | Gammaproteobacteria; Competibacteraceae | I | 2.5 | 458 | 57.7 | 52.1 | 1.4 |
| KWL-0287 | Gammaproteobacteria; Competibacteraceae | I | 1.8 | 428 | 59.2 | 53.5 | 0.0 |
| KWL-0312 | Gammaproteobacteria; Thiobacillaceae | I | 2.7 | 427 | 60.1 | 87.3 | 2.8 |
| KWL-0195 | Gammaproteobacteria; Casimicrobiaceae | I | 1.8 | 342 | 65.2 | 52.1 | 8.5 |
| KWL-0046 | Gammaproteobacteria; Rhodocyclaceae | I | 1.8 | 69 | 64.7 | 70.4 | 0.0 |
| KWL-0056 | Gammaproteobacteria; Rhodocyclaceae | I | 2.7 | 65 | 63.4 | 50.7 | 0.0 |

[1] Obtained with *GTDB-Tk* v1.5.0 based on the GTDB r202.

[2] Obtained with *anvi'o* v7.1 based on the presence of 71 bacterial and 76 archaeal single-copy genes.

**Table S2 (separate tab-delimited file).** Annotation of the coding sequences (CDSs) of the metagenome-assembled genome (MAG) KWL-0264 (*Candidatus* Lamibacter sapmiensis).

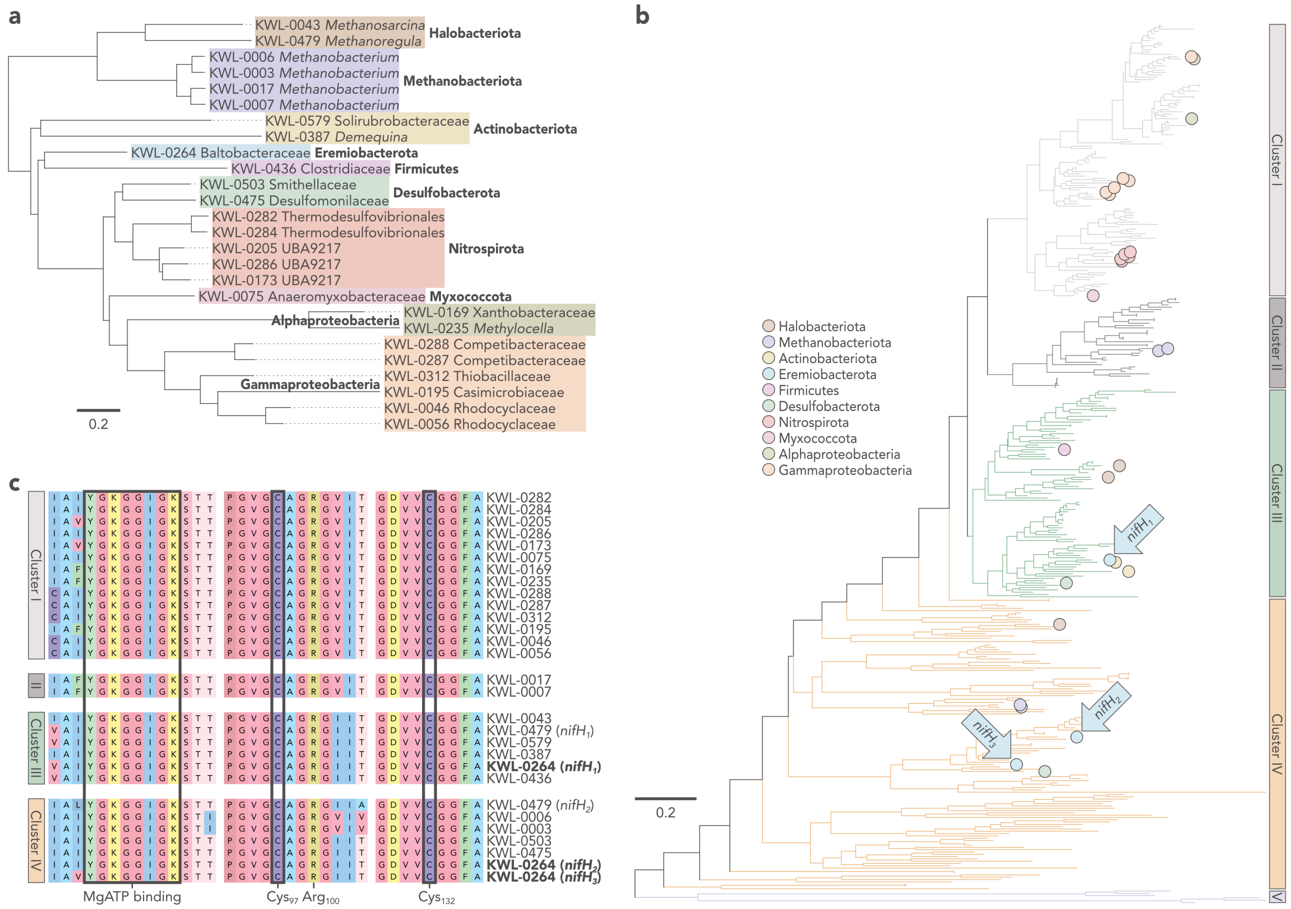


**Fig. S1. a)** Phylogenomic analysis of 26 metagenome-assembled genomes (MAGs) containing *nifH* homologs recovered from tundra soils in Kilpisjärvi, northern Finland. Reduced version of a maximum likelihood tree generated by *GTDB-Tk* v1.5.0 based on the GTDB r202. **b)** Phylogenetic analysis of *nifH* sequences from the 26 MAGs. Maximum-likelihood tree based on the LG+R10 model, rooted at midpoint. The three *nifH* homologs of the Eremiobacterota MAG KWL-0264 are highlighted. **c)** Partial alignment of the *nifH* homologs from the 26 MAGs showing conserved residues associated with the nitrogenase activity. The three *nifH* homologs of the Eremiobacterota MAG KWL-0264 are highlighted.

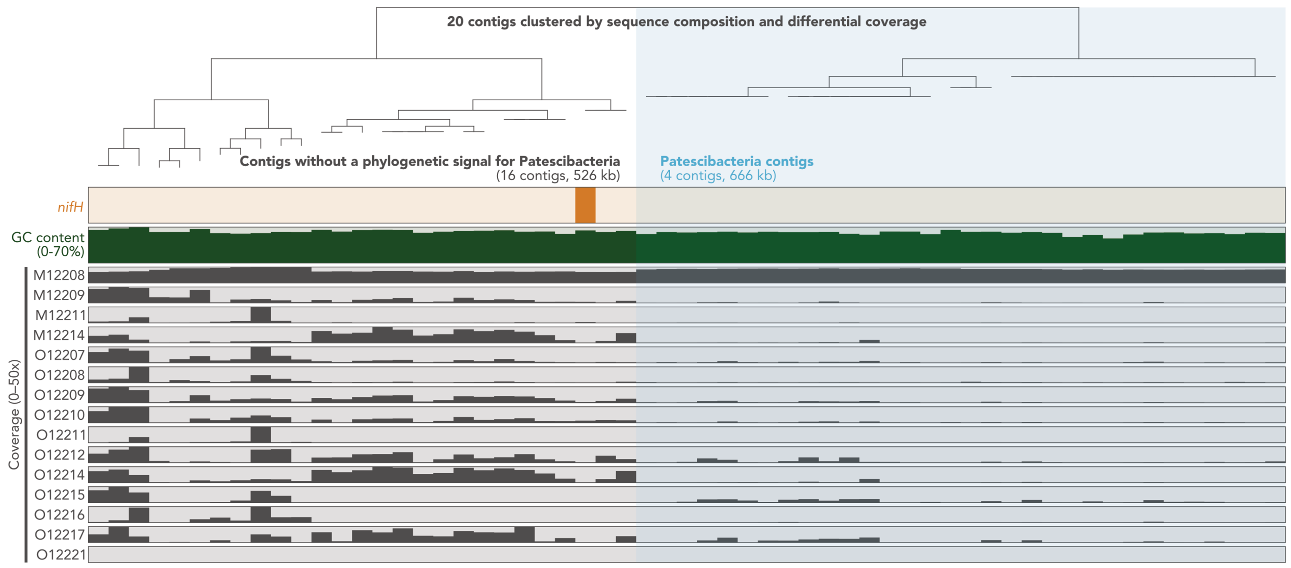


**Fig. S2.** Representation of the 20 contigs of the metagenome-assembled genome (MAG) KWL-0212 showing the location of the *nifH* gene, mean GC content, and mean coverage across 15 fen samples from Kilpisjärvi, northern Finland. For better visualization, contigs ≥20 kb are split into multiple leaves.

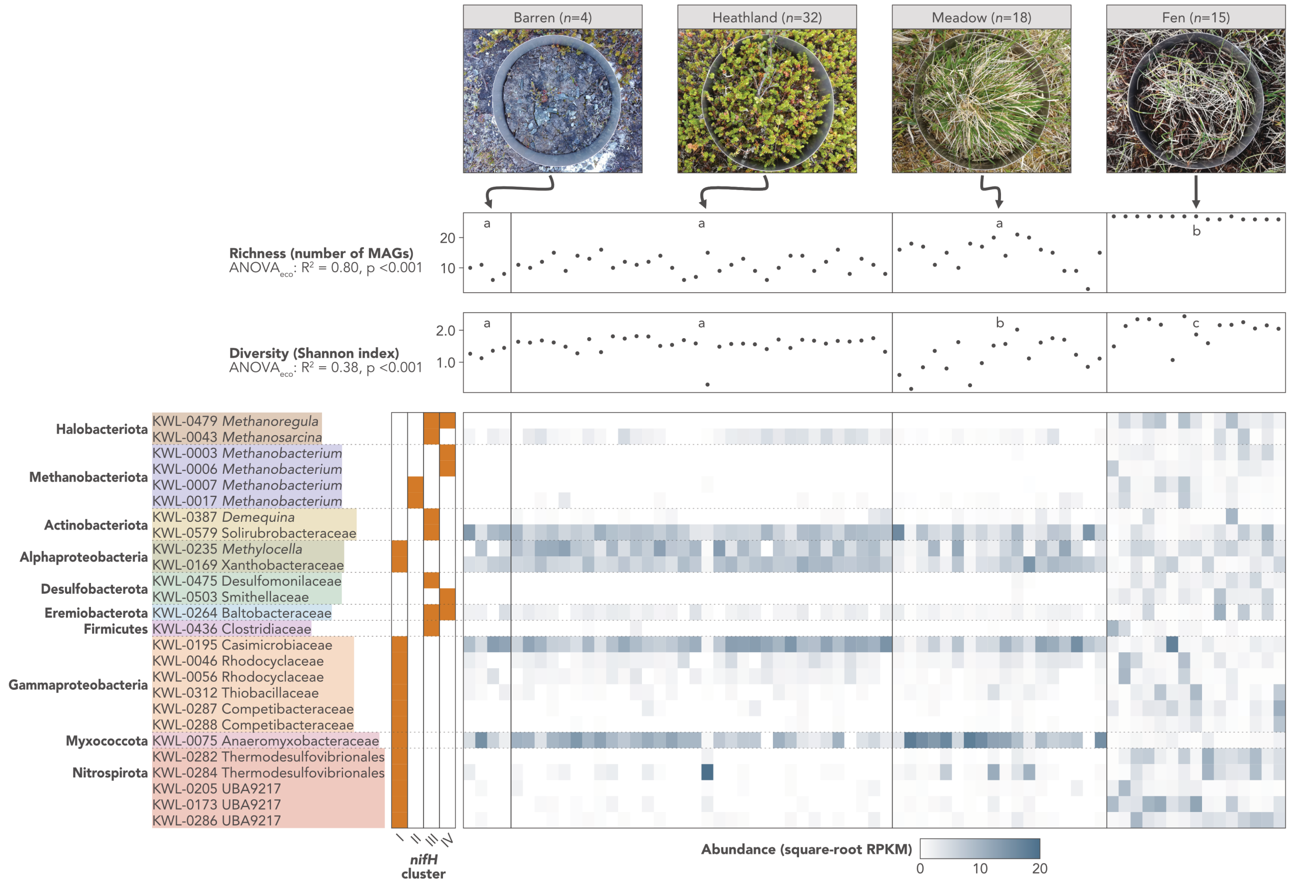


**Fig. S3.** Distribution of 26 metagenome-assembled genomes (MAGs) containing *nifH* homologs across tundra soils in Kilpisjärvi, northern Finland.

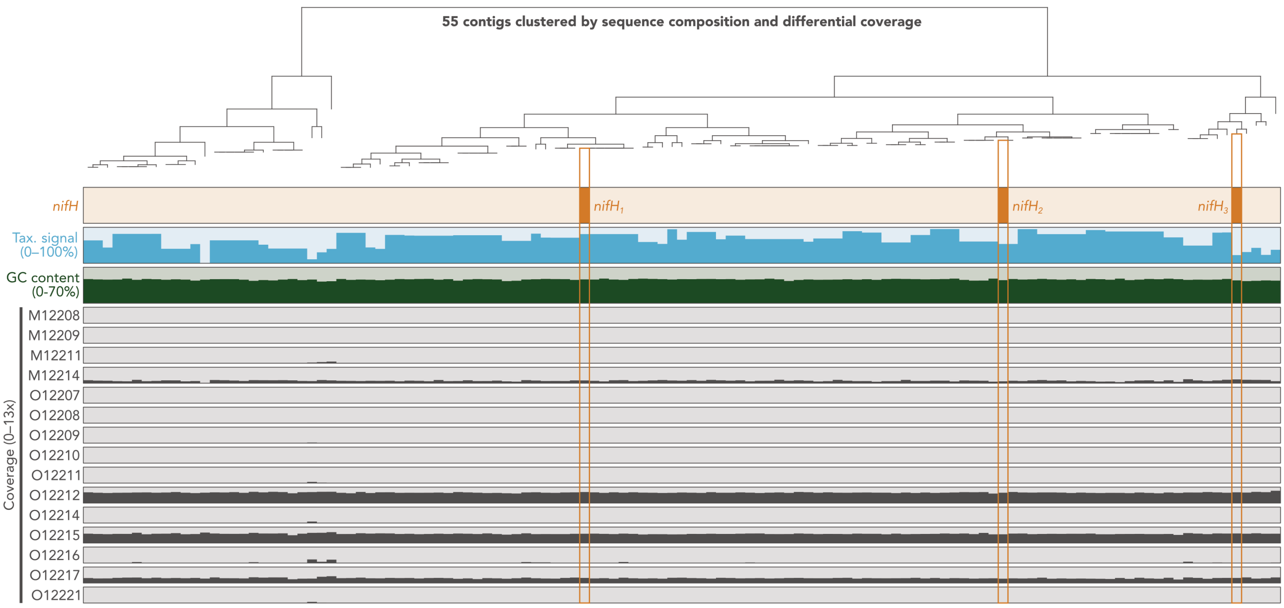


**Fig. S4.** Representation of the 55 contigs of the metagenome-assembled genome (MAG) KWL-0264 (*Candidatus* Lamibacter sapmiensis) showing the location of *nifH* genes, taxonomic signal for Eremiobacterota, mean GC content, and mean coverage across 15 fen samples from Kilpisjärvi, northern Finland. Taxonomic signal represents the proportion of genes in each contig that had the best match with another Eremiobacterota sequence in the GenBank *nr* database. For better visualization, contigs ≥20 kb are split into multiple leaves.

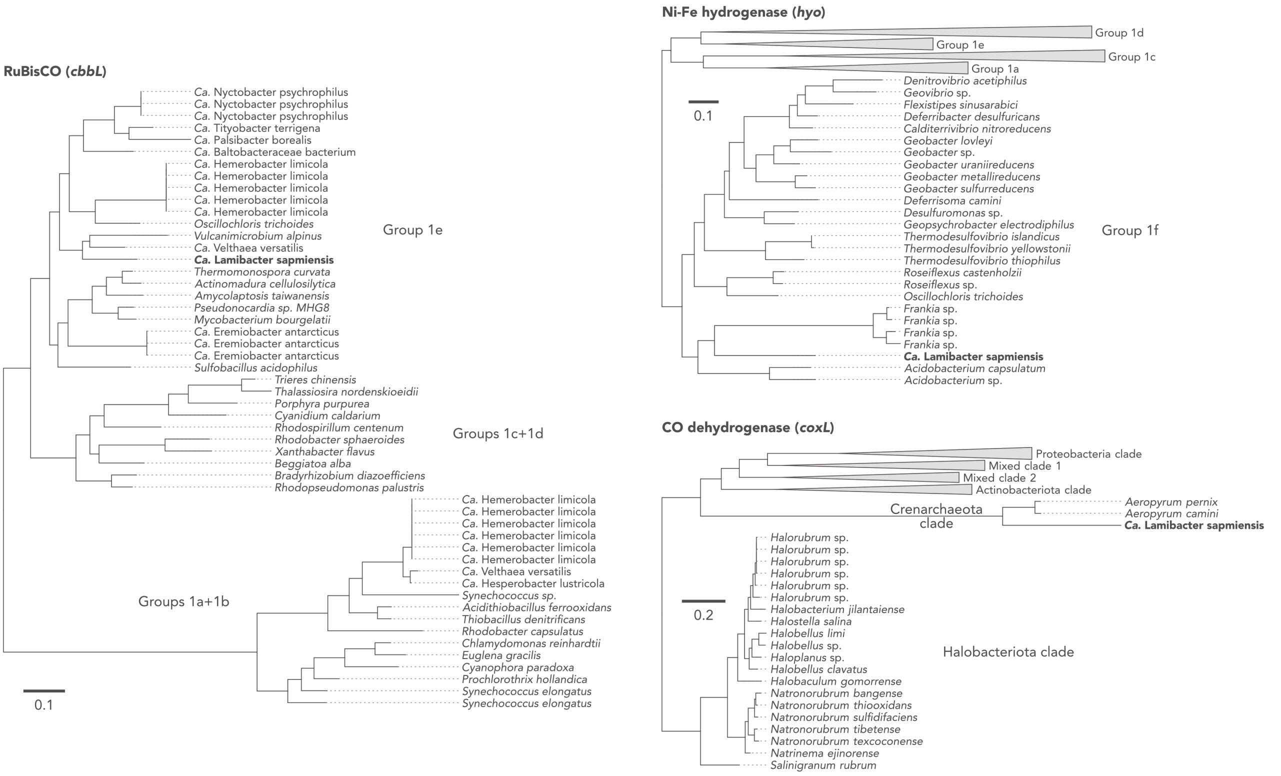


**Fig. S5.** Phylogeny of key genes involved in atmospheric chemosynthesis in the metagenome-assembled genome (MAG) KWL-0264 (*Candidatus* Lamibacter sapmiensis).
